# Unexpected opposing biological effect of genetic risk factors for Parkinson’s disease

**DOI:** 10.1101/702340

**Authors:** Marcus Keatinge, Matthew E. Gegg, Lisa Watson, Heather Mortiboys, Nan Li, Mark Dunning, Deepak Ailani, Hai Bui, Astrid van Rens, Dirk J. Lefeber, Ryan B. MacDonald, Anthony H.V. Schapira, Oliver Bandmann

## Abstract

The additive effect of genetic risk variants on overall disease risk is a plausible but frequently unproven hypothesis. To test this hypothesis, we assessed the biological effect of combined glucocerebrosidase (GCase) and acid sphingomyelinase (ASM) deficiency. Variants in both *glucocerebrosidase1* (*GBA1*) and *sphingomyelinase* (*SMPD1*) are genetic risk factors for Parkinson’s disease. Unexpectedly, ASM deficiency resulted in normalized behaviour and prolonged survival in *gba1*^−/−^;*smpd1*^−/−^ double-mutant zebrafish compared to *gba1*^−/−^. RNAseq-based pathway analysis confirmed a profound rescue of neuronal function and intracellular homeostasis. We identified complete reciprocal rescue of mitochondrial respiratory chain function and abolished lipid membrane oxidation in *gba1*^−/−^;*smpd1*^−/−^ compared to *gba1*^−/−^ or *smpd1*^−/−^ as the underlying rescue mechanism. Complementing *in vitro* experiments demonstrated an unexpected reduction of α-synuclein levels in human cell lines with combined GCase and ASM deficiency. Our study highlights the importance of functional validation for any putative interactions between genetic risk factors and their overall effect on disease-relevant mechanisms rather than readily assuming an additive effect.

**Summary:** The additive effect of genetic risk variants on disease risk is a popular but typically unproven hypothesis. We investigated this hypothesis *mechanistically* for Parkinson’s disease risk factors and provide evidence of an unexpected *rescue* effect on neuronal function and survival.

## Introduction

Genome-wide association studies (GWAS) have become a widely used, powerful tool to identify genetic risk variants for common disorders. However, the overall contribution of each identified genetic risk factor is typically only very small. A popular but largely unproven hypothesis is an additive effect of risk haplotypes, based on the assumption that the risk for a particular disease correlates with the number of risk variants present in an individual. This hypothesis is particularly plausible for variants in genes which encode functionally related proteins.

We tested this hypothesis in the context of genetic risk factors for Parkinson’s disease (PD). GWAS have now identified at least 90 genetic risk factors for PD.^1^ There is particularly strong evidence of an excessive burden of lysosomal disease (LSD) gene variants.^2^ Heterozygous *glucocerebrosidase 1* (*GBA1*) mutations are the most common and strongest genetic risk factor for sporadic PD with a prevalence of ~ 5-20%, depending on the population investigated.^3–5^ Severe *GBA1* mutations confer a considerably higher risk of PD than mild mutations.^6^ However, no additional factors other than age or the severity of the *GBA1* mutation have been identified which would explain the highly variable penetrance of PD in *GBA1* mutations carriers. There is also particularly strong evidence for an association of PD with variants in the *SMPD1* gene (encoding for acid sphingomyelinase (ASM).^7^ Both *GBA1* and *SMPD1* encode lysosomal enzymes that converge on ceramide metabolism. Therefore, we hypothesised that *SMPD1* mutations could modify GBA1 deficiency and in doing so, explain at last a proportion of *GBA1* linked PD.

Zebrafish (*Danio rerio*) are an ideally suited vertebrate model to study gene-gene interactions due to their short generation time, high fecundity and ease of housing. We had previously characterised a *gba1* mutant zebrafish line (*gba1*^−/−^),^8^ and demonstrated its usefulness to study gene-gene interactions.^9^ *gba1*^−/−^ zebrafish faithfully model key features of GCase deficiency, including Gaucher cell accumulation, marked inflammation with microglial infiltration, mitochondrial dysfunction and neurodegeneration.^8^ *gba1*^−/−^ larvae develop normally but *gba1*^−/−^ juvenile zebrafish then rapidly deteriorate from 10-12 weeks onwards and die between 12 and 14 weeks. As both GCase and ASM are lysosomal hydrolases linked to sporadic PD, we hypothesised that additional ASM deficiency could enhance the phenotype and accelerate disease progression in *gba1*^−/−^ zebrafish. As expected, combined GCase and ASM deficiency acted synergistically on key sphingolipid metabolites in the *gba1*^−/−^;*smpd1*^−/−^ double-mutant zebrafish. However, instead of a worsening of phenotypes, we unexpectedly observed markedly prolonged survival and normalisation of the motor phenotype in *gba1*^−/−^;*smpd*^−/−^ zebrafish compared to the *gba1*^−/−^ (single) mutant zebrafish. RNAseq-based pathway analysis confirmed restoration of neuronal health in *gba1*^−/−^;*smpd1*^−/−^ compared to the *gba1*^−/−^. Mechanistic experiments identified a complete reciprocal rescue effect of combined GCase and ASM deficiency on mitochondrial respiratory chain function in *gba1*^−/−^;*smpd1*^−/−^ compared to the marked but distinct mitochondrial dysfunction in *gba1*^−/−^ or *smpd1*^−/−^ single mutant zebrafish. The mitochondrial rescue led to an abrogation of oxidative membrane damage, further reflecting the overall restorative effect of ASM deficiency on neuronal health in GCase deficiency. Complementary *in vitro* work established that the observed mitochondrial rescue effect is autophagy-independent. We also observed an unexpected reduction of α-synuclein levels in human SH-SY5Y cell lines with combined GCase and ASM deficiency rather than the predicted further increase compared to GCase deficient cell lines. Our study highlights the need of functional, mechanistic validation of any putative risk gene interaction for human diseases in suitable model systems rather than readily assuming an additive effect.

## Materials and Methods

### Zebrafish husbandry

All larval and adult zebrafish were housed at the University of Sheffield; experimental procedures being in accordance UK Home Office Animals (Scientific Procedures) Act 1986 (Project license PPL 70/8437, held by Dr Oliver Bandmann). Adult zebrafish were housed at a density of 20 per tank, on a cycle of 14 h of light, 10 h of dark. Adults and embryos were kept at constant temperature of 28°C.

### Mutant line generation and line maintenance

*The gba1*^−/−^ mutant lines was generated using TALEN technology.^8^ The *smpd1*^−/−^ mutant line was generated by the Crispr/Cas9 method as previously described.^10^ The following ultramer template was used: 5’-AAAGCACCGACTCGGTGCCACTTTTTCAAGTTGATAACGGACTAGCCTTATTTTAACTTGCTATTTCTAGCTCTAAAACGGATTGAGGCTTGTGTCTCcCTATAGTGAGTCGTATTACGC). The *smpd1*^−/−^ line was genotyped using primers F 5’-AGCCGTGGTGGTTTCTACAG and R 5’-CCTTCTCTCCCTTGTTCTCG. The *smpd1*^−/−^ line was crossed to *gba1*^+/−^ to generate double heterozygous individuals. These were subsequently in-crossed to generate double mutants. At each in-cross larvae were genotyped at 3dpf by larval tail biopsy as previously described.^11^ Each genotyped was raised in genotype-specific tanks at a density of 10-15 fish per tank. All individuals were re-genotyped at 10 weeks post-fertilisation. *smpd1*^+/−^ were crossed with *gba1*^+/−^ to generate *gba1*^+/−^;*smpd1*^+/−^. These were subsequently in-crossed to generate double mutants, single mutants and WT controls.

### Biochemical activity assays and mass spectrometry

Sphingomyelinase activity was determined using homogenates prepared as follows. Tubes containing twenty embryos (5 dpf) were sonicated in 500 μl MilliQ and centrifuged (11000 rpm). ASM activities were measured using 20 μL supernatant and incubated with substrate HMU-PC (6-hexadecanoylamino-4-methylumbelliferyl-phosphorylcholine 0.66 mM, 20 μL, Moscerdam Substrates, NL) at pH 5.2 and 37 °C during 2 hours. Fluorescence intensity was measured at 415 (ex) and 460 (em) nm using a plate reader (LS55, Perkin Elmer). Lysosomal and mitochondrial enzyme assays as well as mass spectrometry were undertaken as previously described.^8^

### Lipid peroxidation assay

Mitochondria were isolated from the bodies of 3 month old zebrafish. Bodies were homogenised in ice cold mitochondrial isolation buffer. The Abcam lipid peroxidation kit (ab118970) fluorimetric assay was used to measure lipid peroxidation according to manufacturer’s instructions. Results were normalised to WT.

### RNA preparation for gene expression analysis

RNA was prepared from brain tissue of 12 weeks old zebrafish. A Trizol based protocol was used to extract RNA from the tissue. Briefly, individual brains were homogenized in 250 μl TRI Reagent (Sigma) and incubated at room temperature before adding 50 μl chloroform (Thermo Fisher). The samples were centrifuged at 13,300 x g and the top aqueous phase was collected and transferred to a separate tube. RNA was precipitated from the aqueous phase by mixing with equal volume of isopropanol (Thermo Fisher) and centrifugation at 13,300 x g. Precipitated RNA was resuspended in DEPC-treated water (Thermo Fisher) and its concentration and quality were quantified using the Nanodrop 1000 Spectrophotometer. 750 ng of high quality total RNA, with an RNA integrity number (RIN) of 9 or above, was used in the preparation of sequencing libraries using the NEB Ultra II Directional RNA Library Prep Kit (NEB catalogue number E7760), following the polyA mRNA workflow. Libraries were individually indexed and pooled for sequencing. Single-end 100bp sequencing was performed on the Illumina HiSeq 2500 platform using Rapid Run mode with V2 chemistry.

### RNA-seq Analysis

Raw sequencing reads were processed using the bcbio workflow system. The quality of the samples was checked using FastQC and multiqc. The salmon tool (v0.9.01) was used to quantify genes from the zebrafish reference transcriptome (Danio_rerio.GRCz11.98.gtf from Ensembl.org).^12^ The salmon files were then imported into R using the tximport Bioconductor package.^13^ Unsupervised clustering and PCA with DESeq2 revealed a batch effect corresponding to sample preparation date.^14^ Differential expression was performed using DESeq2 incorporating a batch factor into the model. The contrast tested was between *gba1*^−/−^, double mutants and WT; a positive log fold-change indicating higher expression in GBA single mutants. The clusterProfiler Bioconductor package was used to identify enriched pathways in up-(adjusted p-value less than 0.05 and log fold-change > 1) and down-regulated genes (adjusted p-value less than 0.05 and log2 fold-change <-1).^15^

### Gene set enrichment analysis

The analysis was performed with Gene Set Enrichment Analysis (GSEA) software version 4.0.3 (https://www.gsea-msigdb.org/gsea/index.jsp). GSEA preranked analysis was used with default settings except for “Collapse/Remap to gene symbols” set to “No_Collapse”. A ranked list used for the analysis was calculated with each gene assigned a score based on the adjusted p value and the log2 fold change. Zebrafish lysosomal and mitochondrial gene sets were prepared by identifying zebrafish homologues of the genes in human genes sets in Molecular Signatures Database (MSigDB) v7.1.

### Cell culture experiments

SH-SY5Y cells were cultured in DMEM:F12 (1:1) supplemented with 10% foetal calf serum, 1 mM sodium pyruvate, non-essential amino acids and antibiotics. Cells were passaged and transfected every 72 hours with 12.5 nM SMPD1 ON-TARGETplus SMARTpool siRNA (Dharmacon) using HiPerfect transfection reagent (Qiagen) in the absence or presence of 10 μM CBE up to 10 days in culture. Antibodies used were: rabbit LAMP2A antibody (abcam ab125068, 1:1000), mouse α-synuclein antibody (abcam ab27766, 1:1000), mouse LAMP1 antibody (Novus Biologicals NPB2-25155, 1:8000). Human α-synuclein in cell culture media was detected using LEGEND MAX ELISA kit (BioLegend). Total cholesterol levels were measured using the Amplex Red Cholesterol Assay Kit (Invitrogen), while the localisation of cholesterol was detected using Filipin III as per manufacturer’s instructions (abcam ab 133116). Fillipin staining was assessed using a Nikon Eclipse Ti_E inverted microscope (excitation 340 nm and emmission 460 nm; 40X magnification). Images were acquired using a Hamamatsu Orca-Flash camera and NIS Elements AR software.

### Statistical analysis

Graphpad prism V.6 software (Graphpad) was used for statistical analysis and all error bars shown denote the mean ± SD of the mean. All experiments were performed in biological triplicate unless otherwise stated. All data were analysed with either T test, two-way ANOVA. Significance in all enzyme activity assays was determined by two way ANOVA with Tukey’s multiple comparison test.

Supplementary online material: Fig S1 Genetic details of the *smpd1* mutation. Fig S2:beta-hexosaminidase, chitotriosidase and beta-galactosidase activity, across all genotypes. Fig S3: top GO terms upregulated in *gba1*^−/−^ but normalised in double mutants. Supplementary table 1 lists the5 most downregulated genes, rescued in double mutants. Supplementary table 2 lists the 5 most upregulated genes in double mutants. Supplementary table 3 lists the top lysosomal genes rescued in double mutants. Supplementary table 4 lists the top mitochondrial genes rescued in double mutants.

## Results

### *smpd1*^−/−^ zebrafish display abolished acid sphingomyelinase activity and marked sphingolipid accumulation

We identified a single *smpd1* orthologue in zebrafish (ENSDARG00000076121) with 59% shared identity to the human *SMPD1* gene at both the DNA and the protein level. CRISPR/Cas9 technology was used to generate a *smpd1* stable mutant line (*smpd1*^−/−^). The selected mutant allele contained a 5bp deletion and 136bp insertion within exon 3, resulting in a frame-shift and the generation of a premature stop codon (Figure 1A and Supplemental Figure S1). Enzymatic activity of ASM in *smpd1*^−/−^ at 5 dpf was reduced by 93% (p=0.006, Figure 1B). The large reduction in ASM enzymatic activity resulted in a significant increase of key glycolipid substrates in the *smpd1*^−/−^ larvae already at 5 dpf (Figure 1*C*).

**Figure 1:**
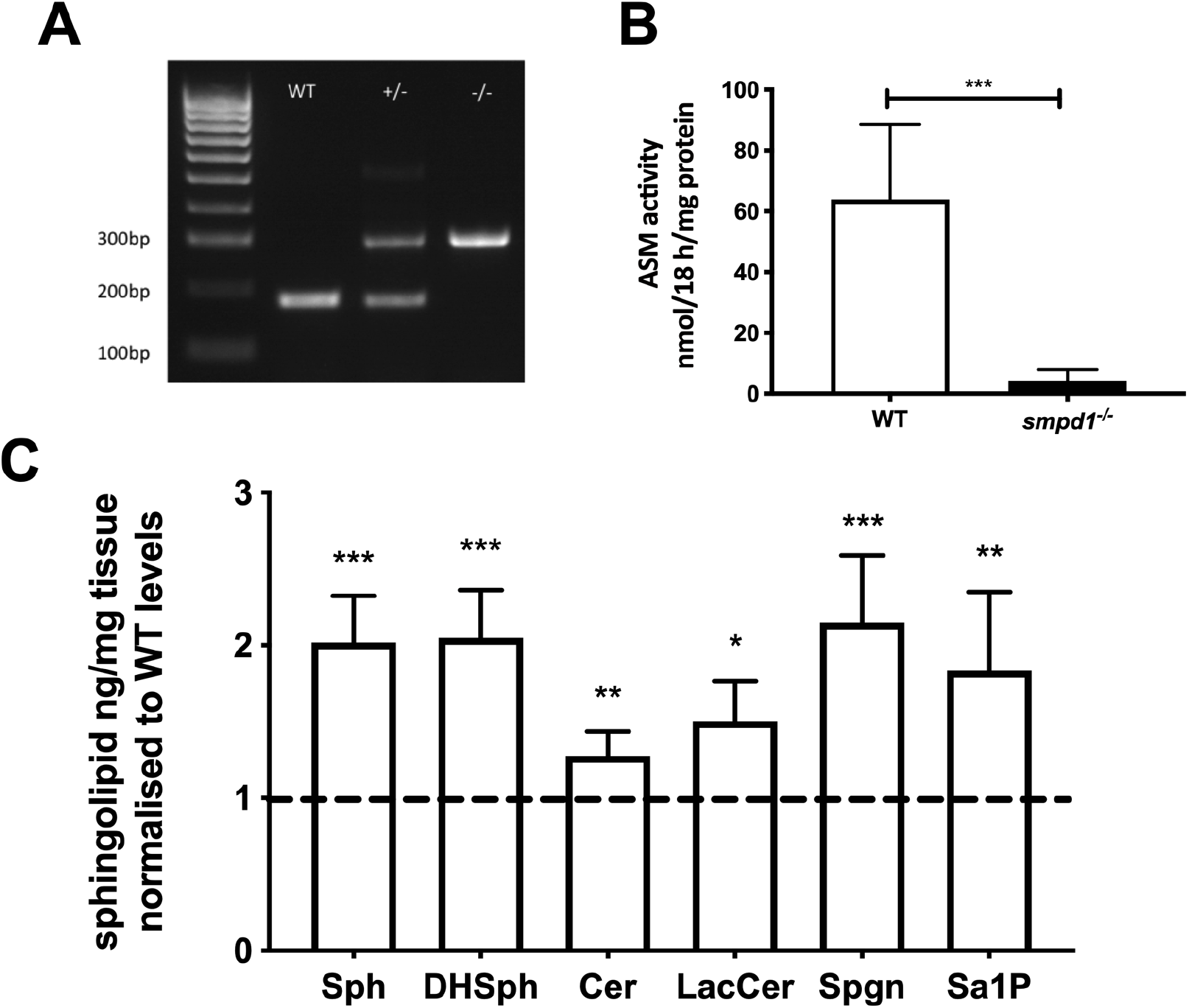
Genetic and biochemical characterisation of the *smpd1*^+/−^ mutant zebrafish line. (*A*) Representative genotyping gel of the *smpd1*^−/−^ alleles (5bp del and 136bp insertion) demonstrating WT, *smpd1*^+/−^ and *smpd1*^−/−^. (*B*) Acid sphingomyelinase (ASM) enzymatic activity compared to WT controls, n=6 larvae at 5 dpf per genotype, p=0.006 by Welch’s t-test. (*C*) Quantification of sphingolipid metabolites, namely Sphingomyelin (Sph), dihydro-sphingomyelin (DHSph), ceramide (Cer), lactosylceramide (LacCer) and Sphinganine 1 phosphate (Sa1P). All metabolites shown are the C18 neuronal species. Data represented are the mean ±SD. ***p<0.001, **p<0.01 and *p<0.05 by two-tailed t-test.

### Combined ASM and GCase deficiency synergistically increases sphingolipid metabolites

We had previously reported marked sphingolipid accumulation in *gba1*^−/−^ zebrafish.^8^ We hypothesised that combined (enzymatic) GCase and ASM deficiency would synergistically increase distinct sphingolipid subtypes. Using mass spectrometry, a comprehensive panel of glycolipid substrates was analysed in the brains of *gba*^−/−^ and *smpd1*^−/−^ single mutant as well as in *gba*^−/−^;*smpd1*^−/−^ double mutant zebrafish and WT controls at 12 weeks of age. A marked additive effect of combined GCase and ASM deficiency was observed for Glucosylceramide levels (the direct substrate of GCase) (Figure 2A). Combined GCase and ASM deficiency also resulted in an additive effect on lactosylceramide, ceramide and sphinganine levels (Figure 2B-D). Sphingosine levels were increased in *gba*^−/−^;*smpd1*^−/−^ compared to WT, reflecting an increase compared to *gba1*^−/−^ but not compared to *smpd1*^−/−^ (Figure 2*E*). Unexpectedly, there was no synergistic effect in sphingomyelin levels in the *gba*^−/−^;*smpd1*^−/−^ double mutants (Figure 2F).

**Fig. 2.**
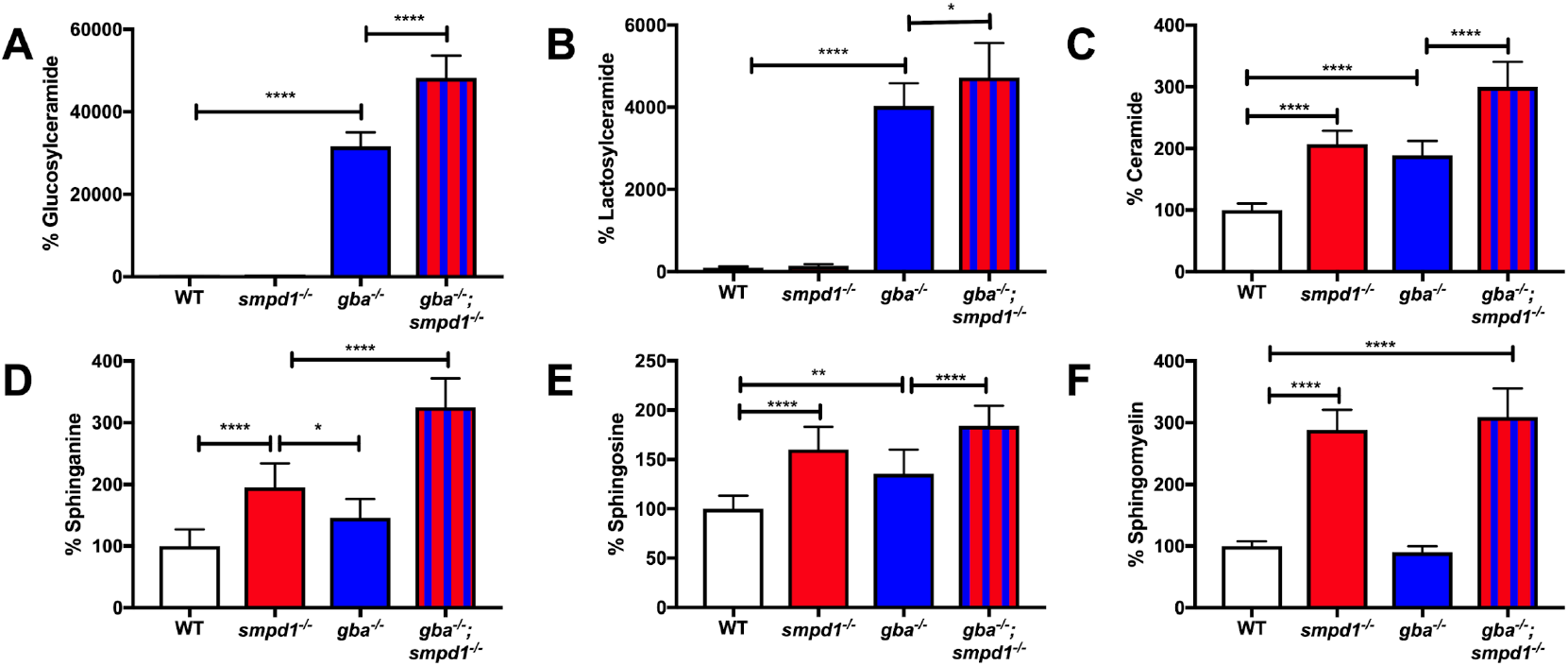
Accumulation of key glycolipids across *gba1*^−/−^ and *smpd1*^−/−^ single mutant and *gba1*^−/−^; *smpd1*^−/−^ double mutant genotypes. (A) Glucosylceramide, (B) Lactosylceramide, (C) Ceramide (D), Sphinganine, (E) Sphingosine and (F) Sphingomyelin levels in WT, single mutant *gba1*^−/−^ vs *smpd1*^−/−^ zebrafish and *gba1*^−/−^; *smpd1*^−/−^ double mutant zebrafish. n=10 of 12 week old zebrafish brains used per group. Data represented are the mean ±SD. ****p<0.0001, ***p<0.001, **p<0.01 and *p<0.05 by 2 way Anova test with Tukey’s multiple comparison.

The inflammation markers chitotriosidase and β-Hexosaminidase are markedly increased in the serum of GD patients and used as biomarkers to monitor disease activity.^16^ We previously observed a marked increase in chitotriosidase and β-hexosaminidase activity in *gba1*^−/−^ zebrafish brain tissue at 12 weeks.^8^ As key GCase substrates were synergistically increased in *gba*^−/−^;*smpd1*^−/−^ double mutant zebrafish, we investigated whether combined GCase and ASM inactivation may also result in a further increase chitotriosidase and β-hexosaminidase activity. Unexpectedly, *gba*^−/−^;*smpd1*^−/−^ double mutant zebrafish displayed a similar increase in chitotriosidase and β-hexosaminidase activity compared to *gba1*^−/−^ (Supplemental Figure 2), suggesting persistent yet unaltered neuroinflammatory states in the double mutants despite a marked synergistic increase in sphingolipid metabolites.

### ASM deficiency unexpectedly prolongs survival in GCase deficiency

The marked additive effect of combined GCase and ASM deficiency on sphingolipid levels led us to hypothesize that ASM deficiency would further worsen the motor phenotype and shorten survival in *gba*^−/−^;*smpd1*^−/−^ double mutant zebrafish. Unexpectedly, genetic inactivation of ASM led to a complete rescue of this behaviour in the *gba*^−/−^;*smpd1*^−/−^ double mutant zebrafish (Supplemental Video S1 (WT), S2 (*smpd1*^−/−^), S3 (*gba1*^−/−^) and S4 (*gba*^−/−^;*smpd1*^−/−^)). Importantly, lifespan was also markedly increased by 22% in *gba1*^−/−^;*smpd1*^−/−^ double mutant zebrafish compared to *gba1*^−/−^ (median survival of 102 dpf in *gba1*^−/−^ and 125dpf in *gba1*^−/−^;*smpd1*^−/−^, p=0.0055, Figure 3A).

**Fig 3.**
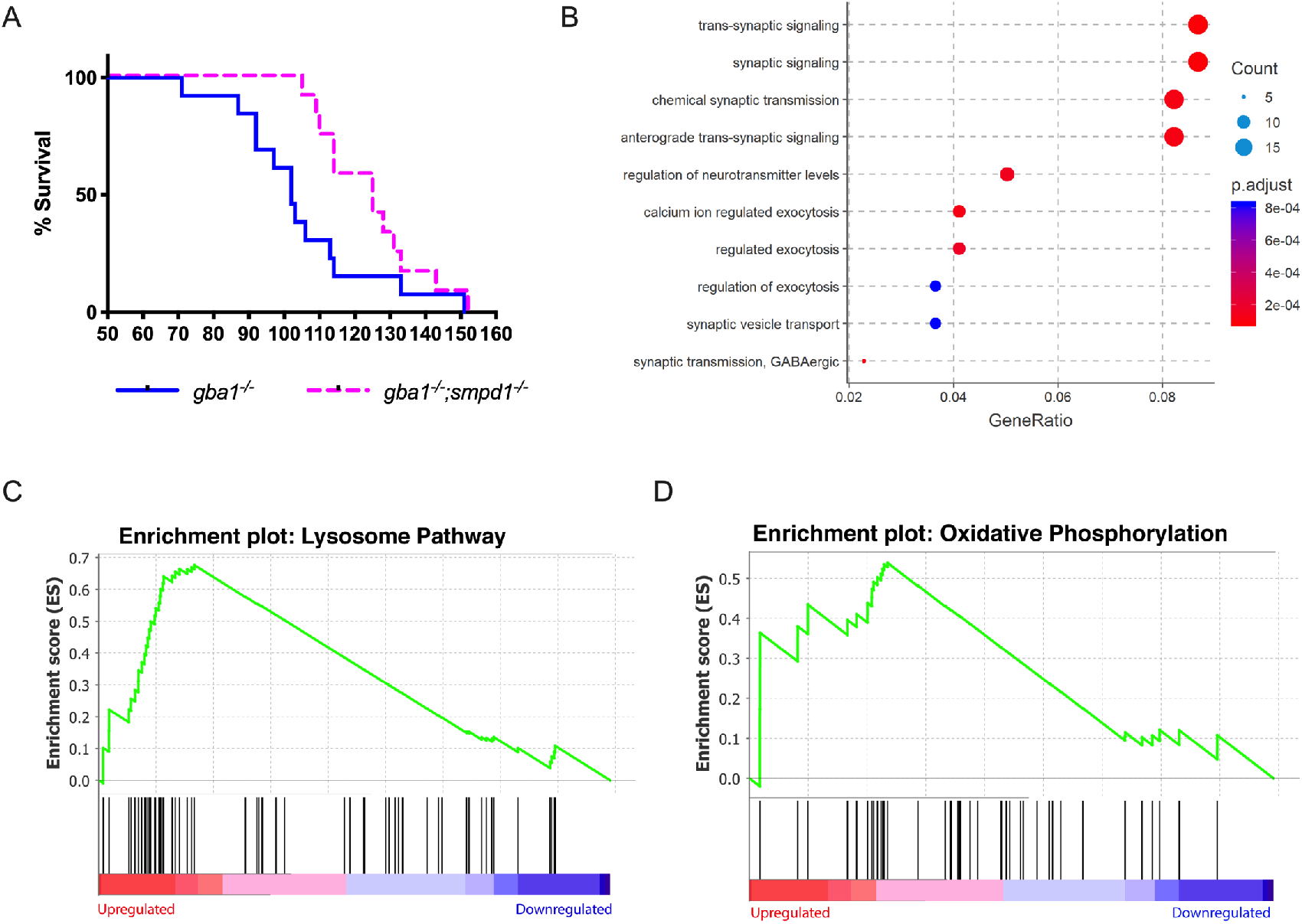
Acid sphingomyelinase deficiency deficiency improves survival and rescues neuronal dysfunction in *gba*^−/−^ zebrafish. (A) Survival analysis of *gba1*^−/−^;*smpd1*^−/−^ double-mutant (n=13) compared to *gba1*^−/−^ single mutant zebrafish (n=12). p= 0.0055 by Gehan-Breslow-Wilcoxon test. (B) The Comparative gene ontology analysis indicates marked global impairment of neuronal function in *gba*^−/−^ but restoration of neuronal health in *gba1*^−/−^;*smpd1*^−/−^. A differential expression analysis was first used to identify genes with statistically significant difference between *gba*^−/−^ and *gba1*^−/−^;*smpd1*^−/−^. Genes with adjusted p-value < 0.05 and log2 fold-change > 1 or <-1 were used to identify enriched biological processes amongst genes over- and under-expressed in *gba*^−/−^. ClusterProfiler identified significant enriched GO (gene ontology) terms within the gene expression changes which were plotted by GeneRatio (ratio of the differentially expressed genes in one particular GO term to the total number of differentially expressed genes). The lists of over- and under-expressed genes were analysed separately. Shown are the ten GO terms with highest gene ratios amongst under-expressed genes; all relating to key aspects of normal neuronal function and homeostasis. These pathways downregulated in *gba1*^−/−^ compared to WT, were normalised in the double mutants. Each GO term is coloured according to the adjusted p-value and ranked according to gene ratio. The size of the point is then scaled according to the number of differentially-expressed genes in the GO term. (C+D) Lysosomal pathway genes and oxidative phosphorylation pathway genes are upregulated in *gba*^−/−^ but normalized in *gba1*^−/−^;*smpd1*^−/−^. The comparison of RNAseq-based transcription levels in the respective pathways between *gba*^−/−^ and *gba1*^−/−^;*smpd1*^−/−^ revealed that both (C) lysosomal pathway genes and (D) oxidative phosphorylation pathway genes were enriched with marked upregulation of both pathways in *gba1*^−/−^ compared to wild-type and *gba1*^−/−^;*smpd1*^−/−^. The x-axis ranks all differentially expressed genes based on the rank metric score from the most upregulated (left) to the most downregulated (right) for either pathway. The vertical black lines show the location of pathway genes in the entire ranked list from the gba1−/− expression changes, compared to WT and double mutants. The y-axis is the degree to which a set of pathway genes is overrepresented at the extremes (up or down-regulated) of the entire ranked list of all differentially expressed genes within the genome. A peak in enrichment score (green line) demonstrates an enrichment of pathway genes amongst all over- or under-represented genes. A sharp peak, demonstrates how highly upregulated each pathway is within the gba1^−/−^ group compared to WT and double mutants.

### RNAseq base pathway analysis confirms restored neuronal health in gba1^−/−^;smpd1^−/−^

We next applied RNAseq-based pathway analysis to further elucidate the underlying mechanisms of the observed rescue effect. The differential gene expression analysis all four genotypes (wild type, *gba1*^−/−^ and *smpd1*^−/−^ single mutants, *gba1*^−/−^;*smpd1*^−/−^ double mutants) identified a total of 512 genes which were dysregulated in *gba1*^−/−^ but rescued in *gba1*^−/−^;*smpd1*^−/−^. Amongst these, there are a downregulation of 219 genes and an upregulation of 293 genes in *gba*^−/−^ compared to wild-type and *gba1*^−/−^;*smpd1*^−/−^ (adjusted p-value ≤0.05, |log2Fold change| ≥1). The five most up- or down-regulated genes are listed in Supplementary Table 1 and Table 2. We next employed ClusterProfiler analysis on gene ontology (GO) categories to identify functionally relevant pathways within the rescued gene sets. Key neuronal pathways including the GO terms for synaptic signalling, chemical synaptic transmission and calcium ion regulated exocytosis were markedly downregulated in in *gba*^−/−^ but normalized in *gba1*^−/−^;*smpd1*^−/−^ (Figure 3B). This suggests that key aspects of neuronal function were restored in the *gba1*^−/−^;*smpd1*^−/−^ double mutants.

We also observed an enrichment of upregulated genes in *gba1*^−/−^ compared to *gba*^−/−^;*smpd1*^−/−^ in a broad range of GO terms, the top 5 of which thought to regulate muscle function. However, since our RNA-seq analysis was carried out on brain tissue, we consider these changes to be of limited relevance only (Suppl Figure 3). Upregulation of the inflammatory signature in *gba1*^−/−^ was retained in the *gba*^−/−^;*smpd1*^−/−^ but not further enhanced (data not shown).

As both GCase and ASM are lysosomal hydrolases, we specifically focused on the effect of isolated GCase deficiency in *gba1*^−/−^ compared to combined GCase and ASM deficiency in *gba1*^−/−^;*smpd1*^−/−^ on lysosome transcriptomic pathways. Gene set enrichment analysis led to the identification of 27 leading-edge, dysregulated lysosomal genes, which account for the pathway’s enrichment signal. The expression of these 27 lysosomal genes was increased in *gba1*^−/−^ compared to wild-type and *gba1*^−/−^;*smpd1*^−/−^ (Fig3C, Supplementary Table 3). Amongst these 27 genes, acid hydrolases contributed the most. Cathepsin L, involved in the initiation of protein degradation, ranked as the top rescued gene. The apparent normalisation of lysosomal gene expression profiles in *gba*^−/−^;*smpd1*^−/−^ was in contrast to the observed marked increase in a wide range of sphingolipid levels in *gba*^−/−^;*smpd1*^−/−^ compared to *gba*^−/−^ *or smpd1*^−/−^ single mutants (see above).

We had previously observed marked mitochondrial dysfunction in *gba1*^−/−^. We therefore also focussed on the analysis of mitochondrial genes involved in the oxidative phosphorylation pathway. This leading-edge mitochondrial gene subset included 16 genes encoding the subunits of the Complex I, II, IV and V in mitochondrial electron transport chain (Supplementary Table 4). Interestingly, gene set enrichment analysis showed an upregulation of this mitochondrial gene subset in *gba1*^−/−^, presumably as a compensatory mechanism to the impaired function of the mitochondrial respiratory chain, but similar mitochondrial gene expression levels in wild-type and *gba1*^−/−^;*smpd1*^−/−^(Figure 6B).

### Reciprocal restoration of mitochondrial function in gba1^−/−^;smpd1^−/−^

We next compared the mitochondrial respiratory chain function across all four genotypes to further determine whether the normalised gene expression levels for oxidative phosphorylation-related genes would be reflected in normalised mitochondrial function. Complex I activity was reduced by 65% in *smpd1*^−/−^ compared to WT levels (p=0.0198, Figure 4A) but restored to 92% of WT levels in *gba1*^−/−^;*smpd1*^−/−^ (p=0.0445, Figure 3A). Complex II was not significantly altered in any of the genotypes (Fig. 4B). Complex III activity in *gba1*^−/−^ was reduced by 45% compared to WT levels (p=0.0091, Figure 4C) as previously observed.^8^ In contrast, Complex III activity in the *gba1*^−/−^;*smpd1*^−/−^ double mutant zebrafish was reduced by only 9% compared to WT levels and thus considerably less pronounced than the reduction observed in the *gba1*^−/−^, suggesting a rescue effect in *gba1*^−/−^;*smpd1*^−/−^. However, this 36% increase in complex III activity in *gba1*^−/−^;*smpd1*^−/−^ compared to *gba1*^−/−^ did not reach statistical significance. Complex IV activity was unchanged in *smpd1*^−/−^ compared to WT, but reduced by 40% in *gba1*^−/−^ compared to WT as previously reported (p=0.0491, Figure 4D). Remarkably, there was a marked improvement of complex IV activity in *gba1*^−/−^;*smpd1*^−/−^ with an increase in activity of 69% compared to *gba1*^−/−^ (p= 0.0005, Figure 4D). Thus, there is reciprocal rescue of mitochondrial respiratory chain function – ASM deficiency normalizes mitochondrial respiratory chain dysfunction in *gba1*^−/−^ and GCase deficiency normalizes mitochondrial respiratory chain function in *smpd1*^−/−^. Malfunction of the mitochondrial respiratory chain can result in oxidative stress and subsequent lipid peroxidation. We therefore investigated next whether the observed rescue in mitochondrial function results in reduced oxidative stress-related damage. Lipid peroxidation was increased in *gba1*^−/−^ brains by 63% above WT levels (p= 0.0214, Figure 4E). As predicted, lipid peroxidation levels were reduced by 70% in *gba1*^−/−^;*smpd1*^−/−^ double mutants compared to *gba1*^−/−^ and thus effectively normalized (p=0.0094, Figure 4E).

**Fig. 4.**
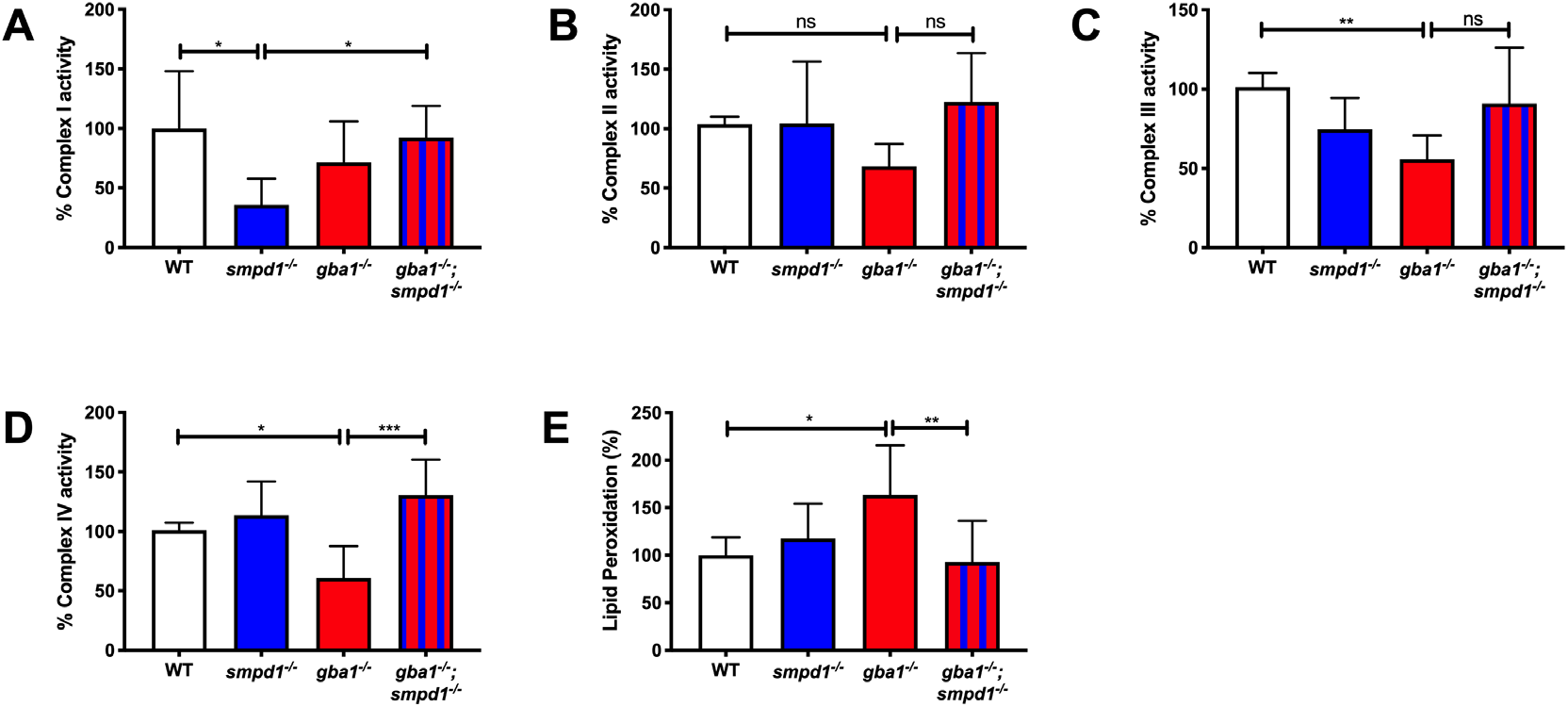
Mitochondrial respiratory chain function and lipid peroxidation. (A) Complex I activity was reduced in *smpd1*^−/−^ by 64±34.77% compared to WT (p=0.0198). Complex I activity was normalized in *gba1*^−/−^;*smpd1*^−/−^ with an increase by 56±21.9% compared to *smpd1*^−/−^ (p=0.0445). (B) Complex II activity was similar across the different genotypes (p>0.05). (C) Complex III activity was reduced in *gba1*^−/−^ compared to WT by 45%±14.99 (p=0.0091) and increased by 35±35.2% in *gba1*^−/−^;*smpd1*^−/−^ compared to *gba1*^−/−^ but this did not reach significance (p>0.05). (D) Complex IV activity was reduced in *gba1*^−/−^ by 40%±26.79 compared to WT (p=0.0491), but completely rescued in *gba1*^−/−^;*smpd1*^−/−^ double mutants with an increase by 69±26.79% compared to *gba1*^−/−^ (p=0.0005). For all mitochondrial complex activity measurements, 6 brains were used for each genotype. (*E*) Lipid peroxidation levels were increased 63±51% in *gba1*^−/−^ compared to WT (p=0.0214), but reduced by 71±43.48% compared to *gba1*^−/−^ and thus effectively normalised to WT levels in *gba*^−/−^;*smpd*^−/−^ double mutants (p=0.0094). For lipid peroxidation experiments, n= 6-8 zebrafish bodies were used for each genotype. Significance in both mitochondrial respiratory chain assays and lipid peroxidation levels was determined by two way ANOVA with Tukey’s multiple comparison test using 12 week brain material. Data represented are the mean ±SD. ***p<0.001, **p<0.01 and *p<0.05.

### The observed mitochondrial rescue effect is autophagy independent

Inhibition of ASM is known to promote autophagy.^17^ We (MG, AS) and others had previously observed impaired autophagy/mitophagy in GCase deficiency.^18,19^ We therefore investigated in our previously established human dopaminergic SH-SY5Y GCase-deficient *in vitro* model whether ASM deficiency may exert its observed rescue effect on mitochondrial dysfunction in GCase deficiency by enhancing autophagy/mitophagy. A siRNA-approach was applied to inactivate *SMPD1/ASM*, whilst GCase was inhibited with conduritol B-epoxide (CBE) for 10 days.^19^ *SMPD1* mRNA levels in the absence or presence of CBE were significantly decreased relative to scrambled (scram) control treated cells (*p* = 0.0003 and 0.0007, Figure 5A). Treatment with CBE alone or in the presence of *SMPD1* siRNA significantly decreased GCase activity (79% and 76% reduction, respectively; *p* < 0.01) relative to scram treated cells. Notably GCase activity was significantly increased in *SMPD1* knock down cells, compared to scram (*p* < 0.05, Figure 5B). Macroautophagy flux was measured by quantifying the levels of LC3-II by western blot, a marker for autophagosome number. Under basal conditions, LC3-II levels were similar in all groups. As expected treatment with bafilomycin A1, which prevents the fusion of lysosomes with autophagosomes, caused an accumulation of autophagosomes, and was similar in all groups (Figure 5C-D). These data suggest that there is neither an impairment nor increase in macroautophagy flux under any conditions. α-synuclein is a key driver of PD pathology.

**Fig. 5.**
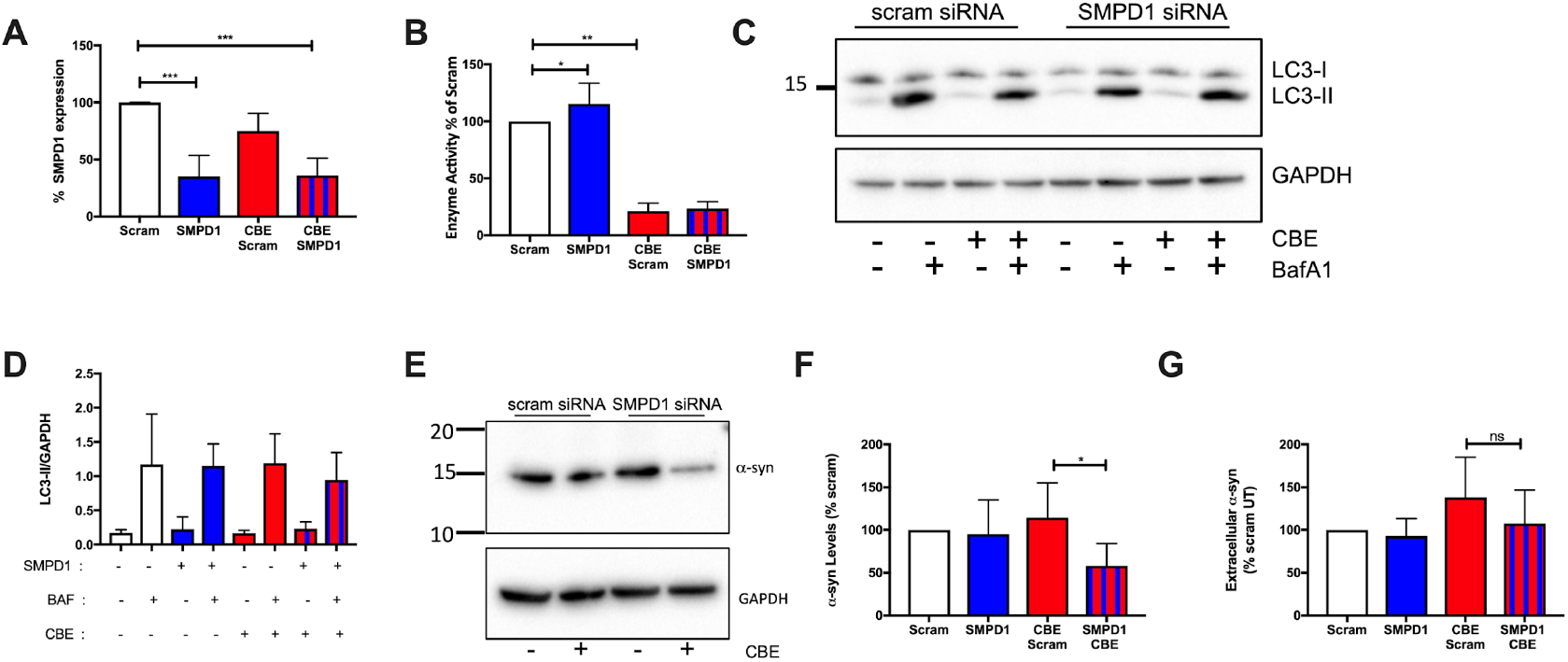
Autophagy in combined GCase and ASM deficiency. (A) *SMPD1* transcript levels after *SMPD1* knockdown (KD) in untreated (n=4) and CBE treated cells (n=3) respectively compared to untreated controls. (B) GCase enzyme activity in *SMPD1* KD compared to Scram control (n=7) and in CBE treated groups compared to Scram untreated control (n=7). (C+D) Macroautophagy flux in *SMPD1* KD cells in the absence and presence of CBE was measured by quantifying LC3-II protein levels using western blotting under basal conditions and following treatment with 100 nM bafilomycin A1 (BafA1). LC3-II protein levels were normalised against GAPDH density and expressed as a ratio, n=3 for all groups. (E+F) Western blotting for TX-100 soluble α-synuclein levels in *SMPD1* KD cells in the absence or presence of CBE. Data were normalised against GAPDH density and expressed as % scram. n = 6 for all groups. (G) Released α-synuclein levels in cell culture media from *SMPD1* KD cells in the absence and presence of CBE for the last 24 hours of treatment, assessed by ELISA. Data were normalised against protein concentration of cells and expressed as % Scram; n = 5. Data represented are the mean ±SD, ***p<0.001, **p<0.01 and *p<0.05 by two way ANOVA with Tukey’s multiple comparison test for all experiments.

### Combined GCase and ASM inhibition results in lowered α-synuclein levels

Given the effect of (isolated) GCase or ASM deficiency on α-synuclein in PD,^7,20^ we also investigated the effect of combined GCase and ASM deficiency on α-synuclein homeostasis. Unexpectedly, the levels of α-synuclein were significantly decreased (rather than increased as predicted) in *SMPD1*-KD + CBE-treated cells (p=0.0139, Figures 5E and F), when compared to CBE-treated cells. The decrease in intracellular α-synuclein levels in *SMPD1-KD*+CBE was not due to increased release of α-synuclein in to the media (Figure 5G) or becoming insoluble (data not shown).

## Discussion

Both *GBA1* and *SMPD1* variants are firmly established inherited risk factors for PD. Additional LSD genes such as *CTSD, SLC17A5* and *ASAH1* have also been implicated.^2^ Biochemically, GCase and acid sphingomyelinase both play a key role in sphingolipid metabolism.^28^ We therefore hypothesized that ASM deficiency would further enhance the phenotype and accelerate disease progression in GCase-deficient zebrafish. However, unexpectedly, we observed a rescue of the “barrel-rolling” phenotype and a marked prolongation of life expectancy, despite clear evidence of an additive effect on the level of the key sphingolipids and their metabolites. The remarkable rescue effect of mitochondrial function in *gba1*^−/−^;*smpd1*^−/−^ on behaviour and survival suggests a central role of mitochondrial dysfunction in GCase deficiency. The profound normalisation of neuronal function in *gba1*^−/−^;*smpd1*^−/−^, as indicated in our RNA-seq based pathway analysis, is in keeping with the observation of the rescued motor phenotype. The remarkable normalisation of intracellular homeostasis is also reflected by the normalisation of both lysosomal and mitochondrial transcriptional pathways.

Partial ASM inhibition restored autophagic dysfunction in AD patient-derived neurons as well as ameliorating the autophagic defect in in AD mice, resulting in a reduction of amyloid-beta deposition and improved memory impairment.^21^ However, in our human dopaminergic cell culture models, macroautophagy flux appeared unaffected in either ASM deficiency alone, or in cells with both decreased ASM and GCase. It should be noted that we had previously observed > 95% GCase inhibition after treatment with 10 uM CBE,^19^ whereas GCase activity was reduced to ~ 80% only in the presence of the transfection reagent required for the knockdown of ASM in this study. This likely explains why the trend for increased α-synuclein levels in the CBE-only treatment arm was not statistically significant but is still comparable to previous results from CBE-mediated GCase inhibition in SH-SY5Y cells^19^ and mouse brain.^22^ The remarkable decrease in soluble monomeric α-synuclein in cells with combined ASM and GCase deficiency may reflect a link between tightly controlled sphingolipid metabolism and α-synuclein homeostasis. Given both enzymes are involved in sphingolipid metabolism, it is probable that changes in the glycosphingolipid profile of the cell contribute to the changes observed. Previous reports have also demonstrated that modulation of the glycosphingolipid profile in GCase deficient cells, other than restoration of GCase activity, can improve α-synuclein homeostasis.^23,24^

The results of our study strongly suggest that experimental confirmation of putative gene-gene interaction will be vital to truly understand the biological role and mechanistic interplay of genetic risk factors for PD and indeed any other human disease.

## Supplemental data

Supplemental data can be found online at …

## Acknowledgements

This work was supported by funding from Parkinson’s UK (G1404 and G1704), the Medical Research Council (MRC, MR/R011354/1 and MR/M006646/1). This research was also supported by the NIHR Sheffield Biomedical Research Centre (BRC).

## Video legends

All videos were taken when the respective zebrafish were 12 weeks of age.

**V-S1:** Normal swimming behaviour in WT. All adults placed into a novel tank swim to the bottom momentarily before resuming standard swimming, balance and buoyancy maintained.

**V-S2:** Normal swimming behaviour in *smpd*^−/−^. All adults placed into a novel tank swim to the bottom momentarily before resuming standard swimming, balance and buoyancy maintained.

**V-S3:** Markedly abnormal swimming behaviour in *gba*^−/−^ with typical “corkscrew” swimming pattern. All adults placed into a novel tank swim in circular motions with balance and buoyancy defects. These increase with frequency and duration at end stage until they need to be culled for humane reasons.

**V-S4:** Normalised swimming behaviour in *gba*^−/−^;*smpd*^−/−^. All adults placed into a novel tank swim to the bottom momentarily before resuming standard swimming, balance and buoyancy maintained.

