## supplementary tables and figures for "Unexpected opposing biological effect of genetic risk factors for Parkinson’s disease" (from another version of this preprint: version 2, 2020-05-09)

**Supplemental Information**

| **Symbol** | **Gene Name** | **Fold Change*** | **adj. p-value** |
| --- | --- | --- | --- |
| slc8a2a | solute carrier family 8 member 2a | 0.014689642 | 2.81E-04 |
| angptl1b | angiopoietin-like 1b | 0.034943137 | 0.003825795 |
| mtnr1c | melatonin receptor 1C | 0.037676856 | 0.012544609 |
| capn8 | calpain 8 | 0.063705159 | 0.019935614 |
| gemin4 | gem nuclear organelle protein 4 | 0.06439142 | 7.02E-04 |

**Supplementary Table 1: The five most downregulated genes rescued by ASM deficiency, ranked according to fold change.** * Fold change was calculated using 2^log2 fold change and represents the ratio of expression level between *gba1^-/-^* and wild-type.

| **Symbol** | **Gene Name** | **Fold Change*** | **adj. p-value** |
| --- | --- | --- | --- |
| LOC100536033 | cathepsin L1-like | 442.3192981 | 0.047451404 |
| myha | myosin, heavy chain a | 232.1802825 | 0.007254479 |
| si:dkey-33i11.4 | si:dkey-33i11.4 | 96.57226654 | 0.003081404 |
| si:dkey-240e12.6 | si:dkey-240e12.6 | 90.5988867 | 0.001662211 |
| pvalb3 | parvalbumin 3 | 74.29246935 | 0.005241277 |

**Supplementary Table 2. The five most 5 upregulated genes rescued by ASM deficiency, ranked according to fold change.** * Fold change was calculated using 2^log2foldchange and represents the ratio of expression level between *gba1^-/-^* and wild-type.

| **Symbol** | **Gene Name** | **Rank Metric Score** |
| --- | --- | --- |
| ctsl.1 | cathepsin L.1 | 8.510105133 |
| cd63 | CD63 molecule | 5.135530472 |
| atp6ap1b | ATPase H+ transporting accessory protein 1b | 5.114915371 |
| lipf | lipase, gastric | 3.06033802 |
| fuca1.2 | alpha-L-fucosidase 1, tandem duplicate 2 | 2.934919357 |
| ctsba | cathepsin Ba | 2.764246225 |
| sftpbb | surfactant protein Bb | 2.624561548 |
| hexa | hexosaminidase A (alpha polypeptide) | 2.622437954 |
| zgc:110239 | zgc:110239 | 2.503591537 |
| acp5a | acid phosphatase 5a, tartrate resistant | 2.363205671 |
| gm2a | GM2 ganglioside activator | 2.312668324 |
| ctsa | cathepsin A | 2.239866734 |
| glb1 | galactosidase, beta 1 | 2.198205709 |
| dnase2b | deoxyribonuclease II beta | 2.142905235 |
| napsa | napsin A aspartic peptidase | 1.98163259 |
| atp6v0a1a | ATPase H+ transporting V0 subunit ca | 1.948094964 |
| ctsh | cathepsin H | 1.844756842 |
| manba | mannosidase, beta A, lysosoma | 1.817932725 |
| galns | galactosamine (N-acetyl)-6-sulfatase | 1.797118545 |
| dnase2 | deoxyribonuclease II, lysosomal | 1.73265183 |
| npc1 | Niemann-Pick disease, type C1 | 1.719917774 |
| gnptg | N-acetylglucosamine-1-phosphate transferase subunit gamma | 1.489329815 |
| man2b1 | mannosidase, alpha, class 2B, member 1 | 1.414416075 |
| ap3b1a | adaptor related protein complex 3 subunit beta 1a | 1.329013348 |
| pla2g15 | phospholipase A2, group XV | 1.1423738 |
| ppt1 | palmitoyl-protein thioesterase 1 (ceroid-lipofuscinosis, neuronal 1, infantile) | 1.051521778 |
| atp6v0ca | ATPase H+ transporting V0 subunit ca | 0.99467504 |

**Supplementary Table 3. The leading-edge gene subset in lysosome pathway**

| **Symbol** | **Gene Name** | **Rank Metric Score** |
| --- | --- | --- |
| atp6ap1b | ATPase H+ transporting accessory protein 1b | 5.114915371 |
| cox6a2 | cytochrome c oxidase subunit 6A2 | 5.065165043 |
| atp5pb | ATP synthase peripheral stalk-membrane | 2.323343754 |
| atp6v0a1a | ATPase H+ transporting V0 subunit a1a | 1.948094964 |
| atp6v0ca | ATPase H+ transporting V0 subunit ca | 0.99467504 |
| ndufb10 | NADH:ubiquinone oxidoreductase subunit B10 | 0.839254677 |
| ndufb6 | NADH:ubiquinone oxidoreductase subunit B | 0.643859327 |
| ndufa11 | NADH:ubiquinone oxidoreductase subunit A11 | 0.643721879 |
| ndufv2 | NADH:ubiquinone oxidoreductase subunit V2 | 0.574424922 |
| cox4i1 | cytochrome c oxidase subunit 4I1 | 0.573632479 |
| sdhc | succinate dehydrogenase complex, subunit C, integral membrane protein | 0.542427838 |
| atp6v1g1 | ATPase H+ transporting V1 subunit G1 | 0.497611314 |
| cox4i2 | cytochrome c oxidase subunit 4I2 | 0.428459734 |
| cox5aa | cytochrome c oxidase subunit 5Aa | 0.396667153 |
| ndufa6 | NADH:ubiquinone oxidoreductase subunit A6 | 0.356948555 |
| ndufc2 | NADH:ubiquinone oxidoreductase subunit C2 | 0.27688098 |

**Supplementary Table 4. The leading-edge gene subset in oxidative phosphorylation pathway.**

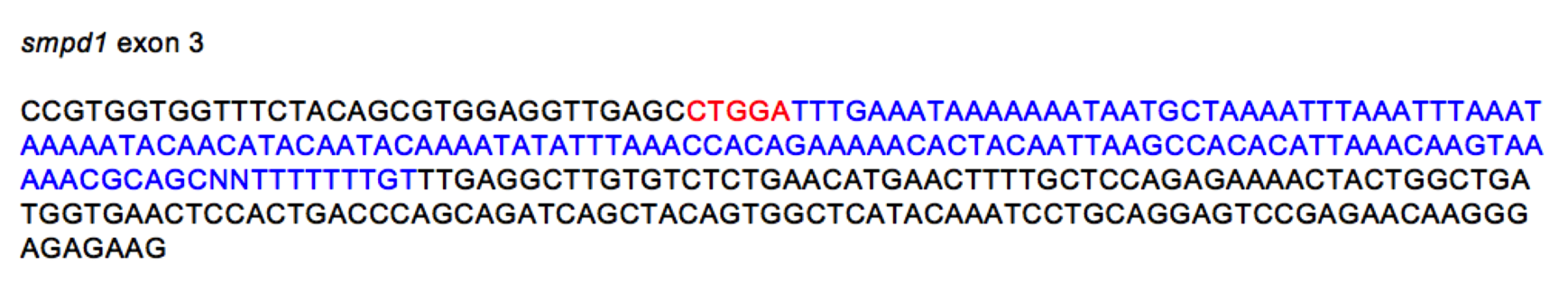

**Fig S1. CRISPR/Cas9 generated *smpd1* allele.**

Using CRISPR/Cas9, we isolated a mutant allele in exon 3 containing a 5bp deletion (red script) with a 136bp insertion (blue script) leading to a frame shift and generation of a premature stop codon at codon 426 (full length WT smpd1: 676 amino acids).

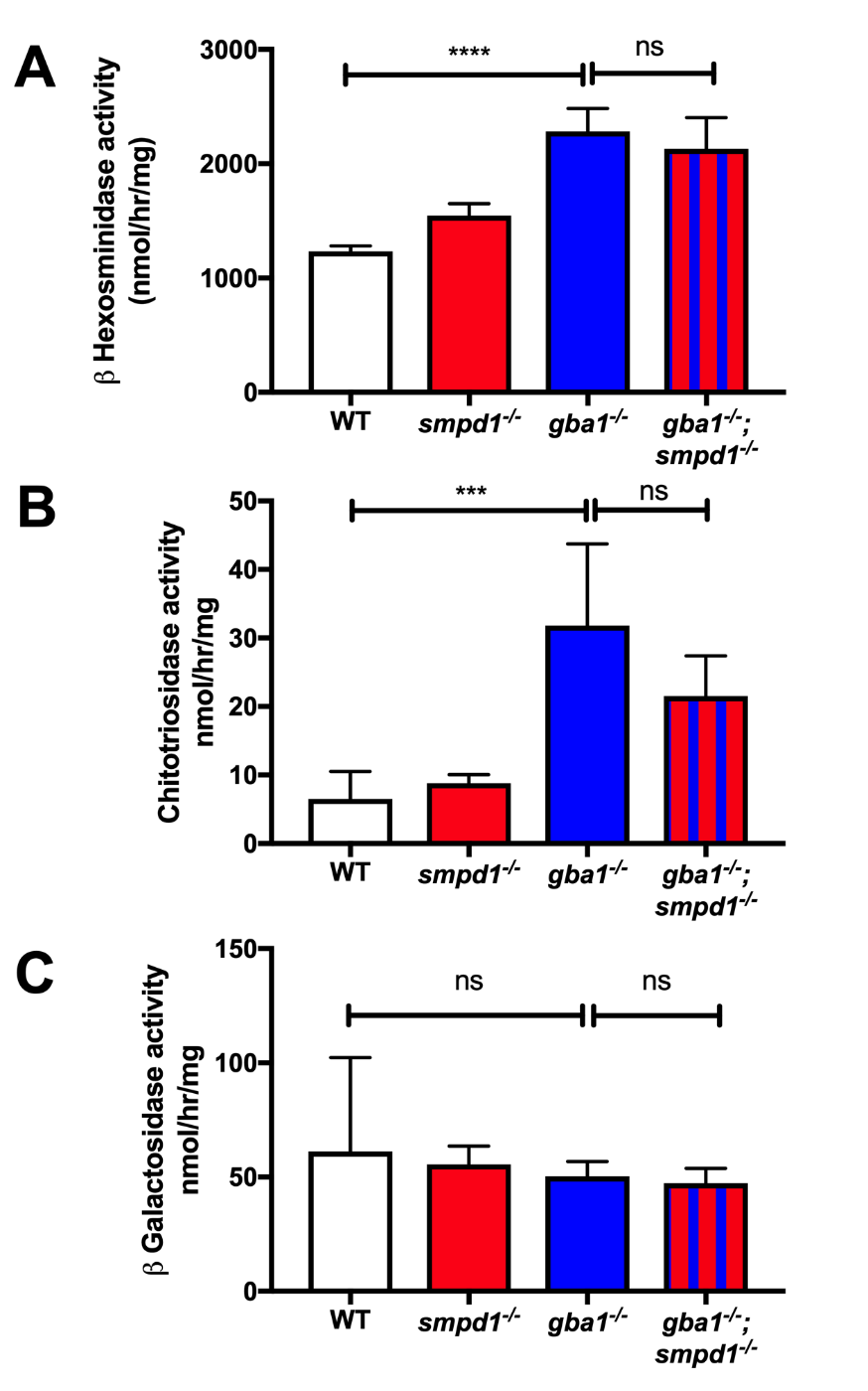

**Figure S2. GD biomarker activation is not substantially altered in *gba1^-/-^;smpd1^-/-^.***

(A) Hexosaminidase activity was not significantly changed between *smpd1^-/-^* and WT, but markedly elevated by 85% in *gba1^-/-^* (2284±200 nmol/hr/mg protein) compared to WT (1234±47.3nmol/hr/mg protein; p<0.0001). *gba1^-/-^;smpd1^-/-^* displayed a similar increase in activity of 70% above WT levels (2131±272.2nmol/hr/mg protein, p<0.0001) compared to *gba1^-/-^*. *(B)* Chitotriosidase activity was not significantly changed in *smpd1^-/-^* compared to WT, but increased in *gba1^-/-^* by 389% (31.8±11.95 nm/hr/mg protein, p=0.0002) compared to WT (6.5±4.0 nm/hr/mh protein) and by 230% in *gba1^-/-^;smpd1^-/-^*showed (21.5±5.1nm/hr/mg protein, p=0.0177) compared to WT. *(C)* Beta galactosidase activity in all genotypes showed comparable activities of approximately 50nm/hr/mg protein. Significance in all enzyme activities assays was determined by two way ANOVA with Tukey’s multiple comparison test with an n of 5 per group using 12 week brain material. Data represented are the mean ±SD. ****p<0.0001, ***p<0.001, **p<0.01 and *p<0.05.

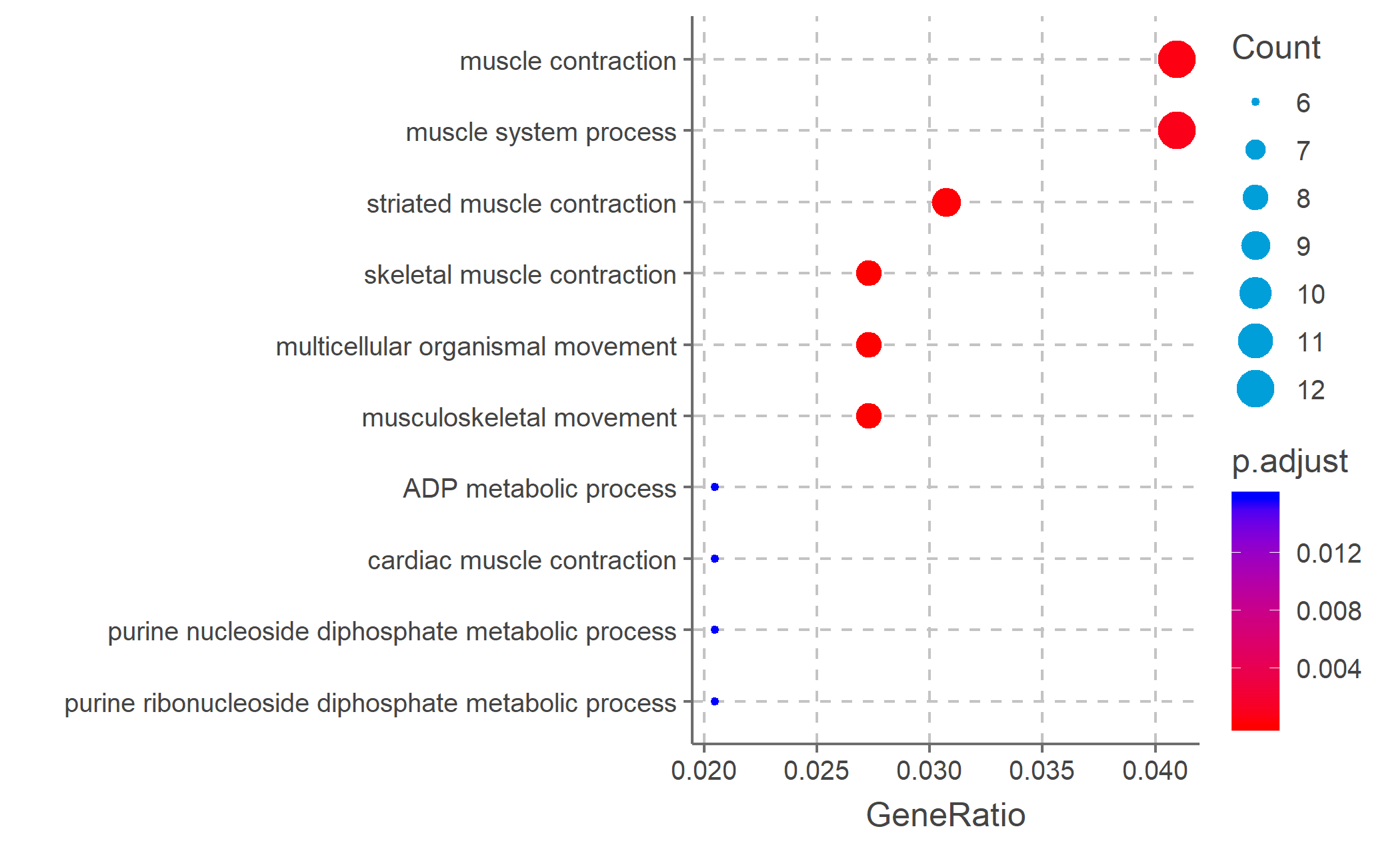

**Figure S3. Over-represented GO terms in the up-regulated genes.**

The top 10 GO terms as identified by ClusterProfiler from the list of statistically-significant genes (adjusted p-value < 0.05 and log2 fold-change > 1 or < -1 respectively. Each point is coloured according to the adjusted p-value for the GO term being over-represented, and the size of the point is scaled according to the number of differentially-expressed genes in the GO term. GO terms are ordered on the y-axis according to the size of the GO-term and adjusted p-value in the case of GO terms with the same number of genes
